# Biochemical characterization of a glycoside hydrolase family 43 β-D-galactofuranosidase from the fungus *Aspergillus niger*

**DOI:** 10.1101/2021.10.27.466152

**Authors:** Gregory S. Bulmer, Fang Wei Yuen, Naimah Begum, Bethan S. Jones, Sabine L. Flitsch, Jolanda M. van Munster

## Abstract

β-D-Galactofuranose (Gal*f*) and its polysaccharides are found in bacteria, fungi and protozoa but do not occur in mammalian tissues, and thus represent a specific target for anti-pathogenic drugs. Understanding the enzymatic degradation of these polysaccharides is therefore of great interest, but the identity of fungal enzymes with exclusively galactofuranosidase activity has so far remained elusive. Here we describe the identification and characterization of a galactofuranosidase from the industrially important fungus *Aspergillus niger*. Phylogenetic analysis of glycoside hydrolase family 43 subfamily 34 (GH43_34) members revealed the occurrence of three distinct clusters and, by comparison with specificities of characterized bacterial members, suggested a basis for prediction of enzyme specificity. Using this rationale, in tandem with molecular docking, we identified a putative β-D-galactofuranosidase from *A. niger* which was recombinantly expressed in *Escherichia coli*. The Gal*f*-specific hydrolase, encoded by *xynD* demonstrates maximum activity at pH 5, 25 °C towards 4-Nitrophenyl-β-galactofuranoside (*p*NP-β-Gal*f*), with a K_m_ of 17.9 ± 1.9 mM and V_max_ of 70.6 ± 5.3 μmol min^−1^. The characterization of this first fungal GH43 galactofuranosidase offers further molecular insight into the degradation of Gal*f*-containing structures and may inform clinical treatments against fungal pathogens.

## Introduction

The 5-membered ring form of galactose, β-D-galactofuranose (Gal*f*) is a key structural component of many pathogens, in which glycans containing the sugar can be highly immunogeneic^1^. Despite its presence spanning fungi, bacteria, protozoa, sponges and green algae^2^ the monosaccharide is not present in mammalian tissue, offering a clear target for therapeutics^1–3^. The occurrence of Gal*f* differs greatly from its pyranoside ring form, which is found widely across mammalian biology^4^. Despite the importance of motives containing this monosaccharide, knowledge is limited regarding enzymes that are active on glycans containing Gal*f*.

In many fungi Gal*f* is a key structural component of the cell wall and is present in secreted molecules, it has been shown to drive immunogenic responses in mammals^5–8^. Gal*f* containing polysaccharides are found in pathogens such as *Mycobacterium tuberculosis*, *Cryptococcus neoformans* and *Aspergillus fumigatus* which combined are responsible for over 2 million deaths worldwide per annum^5,9,10^. Gal*f* can constitute either the core or the branched sections of polysaccharides. In *M. tuberculosis* arabinogalactan (AG), a core of ~35 Gal*f* moieties with alternating β-1,5 and β-1,6 linkages is decorated with a variety of arabinose branches (Fig. 1A)^11–13^. *Mycobacterium* knockouts of the gene encoding UDP-galactopyranose mutase are unable to produce Gal*f* and fail to proliferate *in vitro*, showing that galactan synthesis is essential for replication of *Mycobacterium*^14^. In Aspergilli the predominant Gal*f* containing structure is galactomannan, a polysaccharide formed by a α-1,2/α-1,6-linked mannose backbone with Gal*f* side chains (Fig. 1B)^15–18^. The detection of this galactomannan is utilised to test for *Aspergillus* in a clinical setting^19^. Galactomannan plays a key role in cell integrity whereby *A. fumigatus* mutants lacking the ability to insert galactomannan into their cell walls suffer severe growth impairment^20^. Additionally Gal*f* is also found in a variety of glycoconjugates: such as O-antigens of *Escherichia coli* lipopolysaccharide^21^, *Klebsiella pneumonia* galactan-I repeating unit^22^, lipophosphoglycans and glycoinositolphospholipids of leishmaniosis-causing protozoa *Leishmania major*^23^ and as part of glycoinositolphospholipids and N- and O-glycans on proteins secreted by *A. niger*^24,25^ (Fig. 1C).

**Figure 1:**
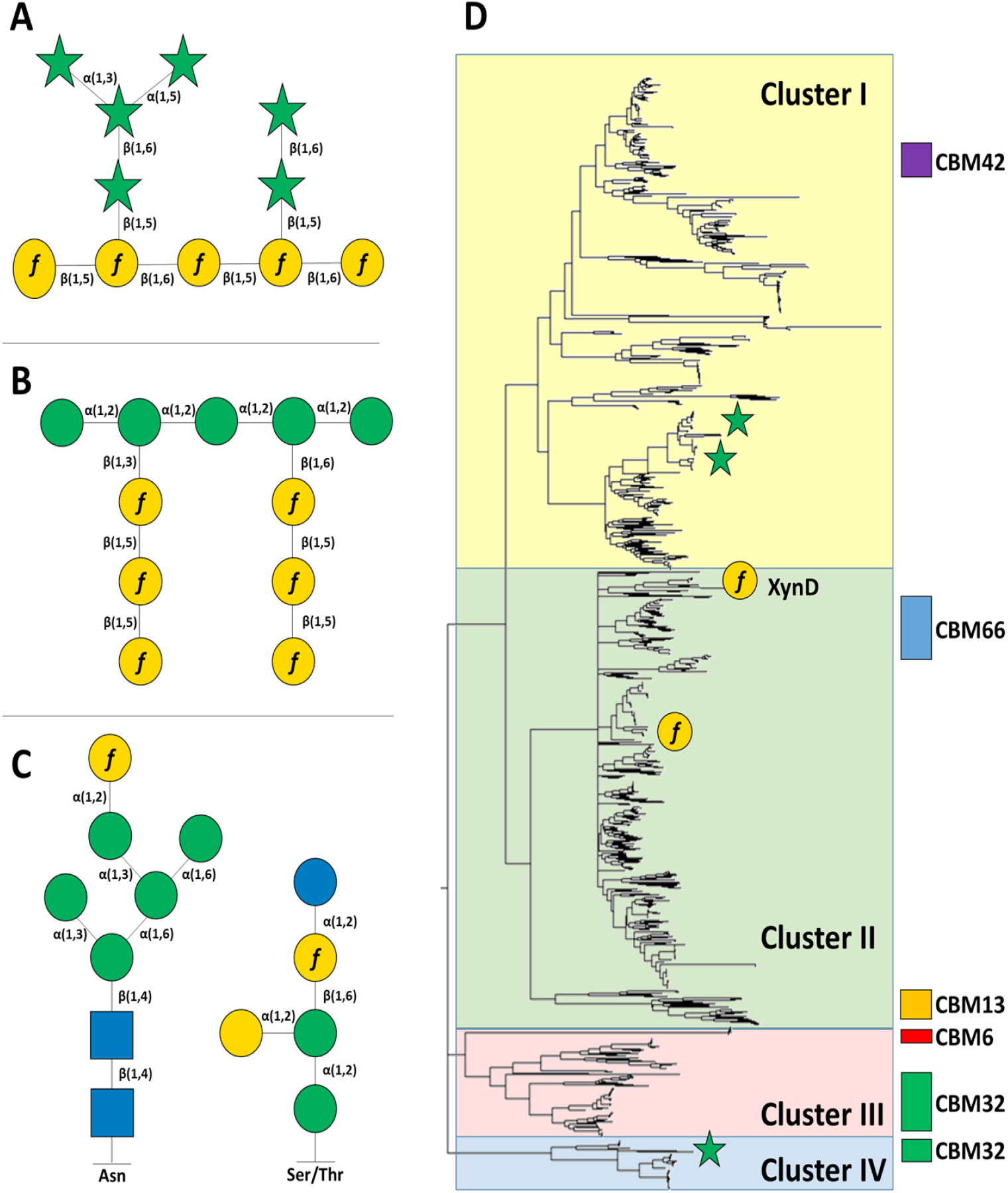
The presence of Gal*f* throughout the fungal and bacterial kingdoms is not uniform and contains a variety of structures of both Gal*f* backbones and Gal*f* side chains, for example. **a** Structure of *M. tuberculosis* arabinogalactan fragment, **b** *A. fumigatus* galactomannan fragment, **c** N- and O-linked glycans. **d** Phylogenetic tree of GH43_34 family proteins with distinct clades visualised. Known activities are demonstrated as follows: β-galactofuranosidase (yellow circle), α-arabinofuranosidase (green star).

In a clinical setting, understanding the turnover of immunogenic sugars is of great importance to treating patients with invasive symptoms from various fungal infections. Additionally, comprehension of the metabolism of Gal*f* and poly-saccharides containing it offer opportunities for industrial application. Fungi such as *A. niger* are often utilised as expression hosts for complex pharmaceuticals with human or other mammalian recipients^26,27^. Therefore considering the metabolism of such immunogenic sugars as Gal*f* is important when designing such processes to avoid potential immune cross-reactivity.

Recently characterized bacterial β-galactofuranosidases (Gal*f*ases) begun to enter the literature, including Gal*f*ases specific for Gal*f* as well as bifunctional enzymes with activity on Gal*f* and Ara*f*^28–30^. However, research into fungal Gal-*f*ases is considerably less advanced. A Gal*f*ase from *A. niger* has previously been employed as a tool for glycoconjugate analysis where proof of concept studies demonstrated the enzyme was able to remove Gal*f* from glycoproteins with *O*-linked (glucoamylase GAM-1) and *N*-linked (α-galactosidase A) Gal*f* containing glycans^31,32^. In addition the supernatant of *A. niger* cultures can hydrolyse the biologically relevant fungal galactomannan^33^. However, at a gene level the source of this activity lacks complete characterization. *A. niger* α-L-arabinofuranosidases AbfA and AbfB from GH families 51 and 54 have a dual activity and can hydrolyse both pNp-α-Ara*f* and pNp-β-Gal*f*^33^ but are not active on fungal galactomannan, indicating other β-galactofuranosidases remain un-characterized. While undertaking the work described here, discovery of GH2 family β-galactofuranosidases GfgA and GfgB in *A. nidulans* has been reported, thus identifying the first specific fungal β-galactofuranosidases^34^. The biological function of Gal*f*ase enzymes *in situ* remain unclear, however, it has been suggested that the breakdown of galactomannan and other Gal*f* epitopes may be utilised as a carbon source, via degradation by Gal*f*ase, during carbon limitation to produce a source of galactose^32^.

One reason for limited understanding of Gal*f*ases stems from the under examined nature of many large GH families. For example, members of GH43-34, comprised to our knowledge only 6 biochemically characterized bacterial proteins at the time of writing, which all showed either β-D-galactofuranosidase or α-ʟ-arabinofuranosidase activity^35–37^. As bi-functionality of Ara*f*ases and Gal*f*ases seems prevalent^28,33,34^ and Ara*f* and Gal*f* are structurally similar we hypothesized that fungal Gal*f*ases were potentially harboured in GH families containing both Ara*f*ases and Gal*f*ases.

Therefore, in this work, we describe how we identified an *A. niger* GH43-34 enzyme, XynD, as a potential Gal*f*ase candidate via a combination of phylogenetic analysis, structural modelling and substrate docking. XynD is known to be expressed during *A. niger* interaction with plant biomass, however the biochemical function of XynD had yet to be elucidated. We express and characterize the enzyme, XynD, and demonstrate that the enzyme is specific for Gal*f*. This Gal*f*ase enzyme represents the first identified and characterized fungal GH43 family galactofuranosidase.

## Results

### Phylogenetic analysis of XynD

We aimed to elucidate the role and biochemical activity of the GH43-34 enzyme from *A. niger*. Members of this sub-family exhibit different biochemical specificities, as either galactofuranosidases, arabinofuranosidases or arabinanases. Therefore, we investigated whether biochemical activity could be inferred from phylogenetic analysis and comparison of characterized enzymes with known activities^35,38,39^ in this subfamily. Catalytic domains of all fungal and bacterial entries from GH43-34 on the CAZy database^40^ were selected for phylogenetic analysis. Construction of a phylogenetic tree via Maximum likelihood analysis revealed four distinct clusters (cluster I-IV) within the family (Fig. 1D). The distribution of sequences over the clusters does not follow the evolutionary relationships between species, with a mixture of fungal and bacterial sequences found in cluster I and II, while cluster IlI and IV comprised only bacterial members. For those organisms containing multiple GH43-34 members, different enzymes can be located in different clusters. Cluster I contains two arabinofuranosidases (ALJ58905.1 and AAO78780.1) whilst cluster II contains one biochemically characterized enzyme, a galactofuranosidase (ALJ48250.1). Based on this separation, we hypothesize that *A. niger* XynD (An11g03120, GenBank: CAK40644.1), which is also found in cluster II, may have galactofuranosidase activity. Cluster IV contained one arabinofuranosidase (AAO78767.1) whilst cluster III contained no characterized members. Additionally through utilising the CAZy database we identified CBM (Carbohydrate Binding Module) protein domains that co-occur with GH43-34 members, namely CBM6, CBM13, CBM32, CBM42 and CBM66 (Fig. 1). See Fig. S1 for bootstrap values and fungal member locations.

### Modelling and substrate docking

To investigate the potential activity of *A. niger* XynD in more detail we created a model of the protein’s structure based on the crystal structure of GH43-34 arabinofuranosidase from *Bacteroides thetaiotaomicron* BT_3675 (PDB 3QZ4)^41^. Of the crystallised GH43 subfamily 34 members, 3QZ4 gave the greatest percentage coverage for modelling (89%), with 37% sequence identity. GH43 family enzymes display a five-bladed β-propeller fold^42^, the presence of this fold was also predicted in the XynD model. The XynD model composes of a monomer 36.4 kDa in size and aligns with one of the monomers of the 3QZ4 dimer macromolecule. No kinetic or substrate binding data is available for 3QZ4. Therefore, using this model, we applied molecular docking simulations to gain insights into the functional binding of potential substrates in the active site. Docking studies showed energetically favourable binding of galactofuranose to the catalytic domain of XynD. Substrate docking suggested five residues involved in catalysis that were energetically favourable: Asp-28, Asp-151, Glu-199, Trp-219 and Arg-286. The first three of these (Fig. 2A) correspond with the catalytic Brønsted base, pKa modulator and catalytic acid that have been identified in GH43 family members^42^. Molecular docking of arabinose in the active site demonstrated that interaction between this sugar and residues Asp-151 and Glu-199 was lacking and thus suggested a specificity for galactofuranose as opposed to arabinofuranosidase or galactofuranosidase/ arabinofuranosidase dual activity. Analysis of the substrate binding site suggests a restricted binding pocket for galactofuranose (Fig. 2B).

**Figure 2:**
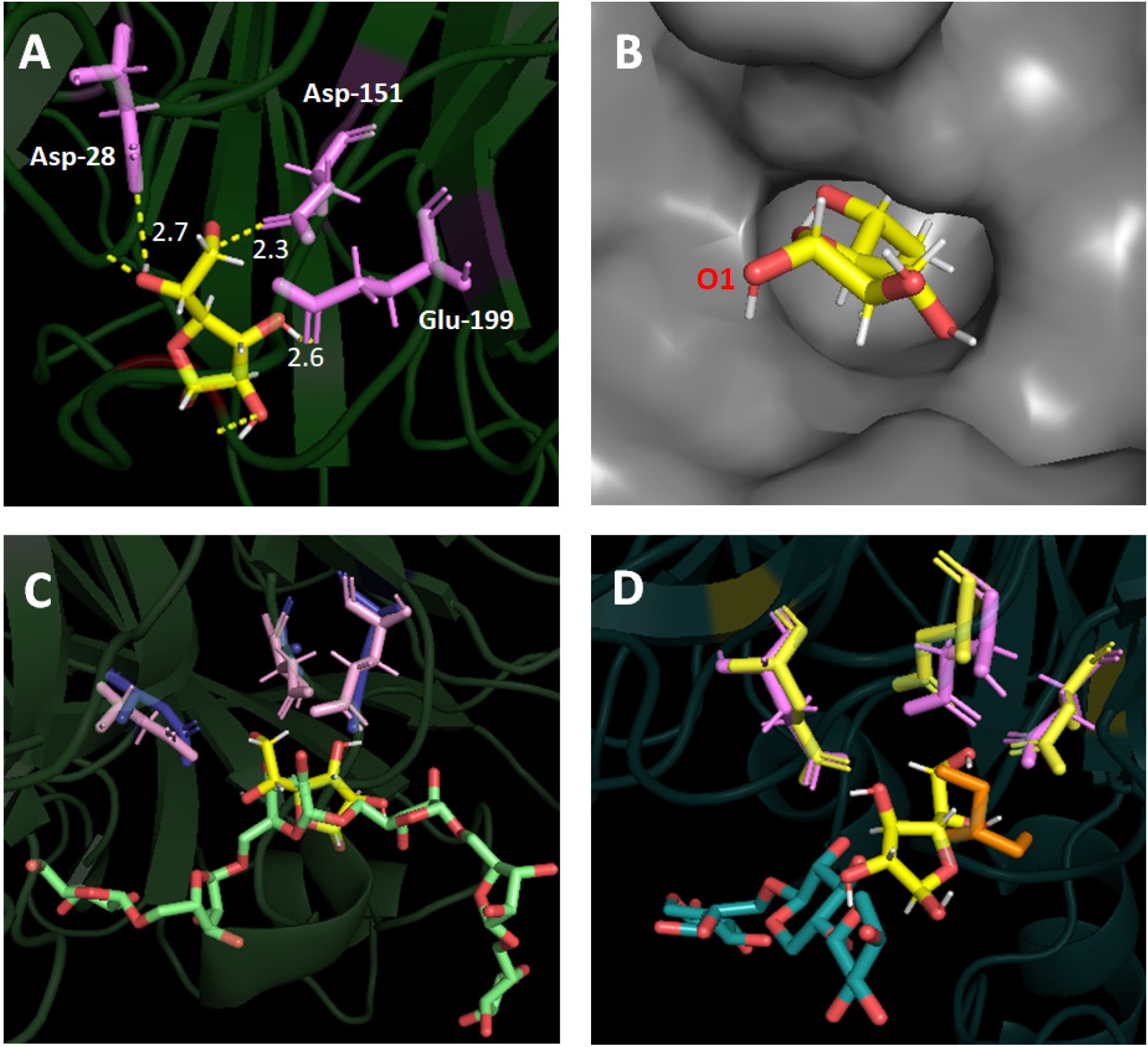
**a** Molecular docking (AutoDock VINA as implemented in YASARA) of galactofuranose (yellow) in a model of *A. niger* XynD generated using PDB 3QZ4 as a template. Catalytic residues Asp-28, Asp-151, Glu-199 are displayed in lilac with bonding lengths indicated in ***Å* b** XynD surface highlighting restricted binding pocket **c** GH43 *C. japonicus* alpha-L-arabinase Abf43A (PDB 1GYE) depicted with catalytic triad (blue) and arabinohexaose ligand (green) overlaid with catalytic triad of XynD (liliac) and Gal*f* ligand (yellow) **d** *B. subtilis* arabinoxylan arabinofuranohydrolase BsAXH-m2,3 (PDB 3C7G) depicted with catalytic triad (yellow), glycerol (orange) and xylotetraose ligand (blue) overlaid with catalytic triad of XynD (liliac) and Gal*f* ligand (yellow).

In order to understand the roles of residues in XynD the enzyme was compared with the crystal structures of related enzymes. Comparison of the XynD model with the *Cellvibrio japonicus* alpha-L-arabinanase Arb43A complexed with arabinohexaose (1GYE) (Fig. 2C) and *Bacillus subtilis* arabinoxylan arabinofuranohydrolase BsAXH-m2,3 in complex with xylotetraose (3C7G) (Fig. 2D), suggested the catalytic triad was conserved. Differences were observed between these proteins in residues involved in hydrophobic stacking. BsAXH-m2,3 contains several residues involved in hydrophobic stacking: Phe-244 at the II subsite, Trp160 at the III (−1) subsite and Trp-101 at the IV (+1) glycerol containing subsite. XynD contains an equivalent residue for Phe-244 (XynD Trp-219) and Trp-101 (XynD Trp-86), but not for Trp-160. Alignment also identified Arg-286 as an equivalent residue to the BsAXH-m2,3 Arg-321 involved in +1 subsite stability. 1GYE shares the three conserved catalytic residues with XynD but the Phe-114 of 1GYE involved in the high binding affinity of the enzyme to arabinan has no corresponding residue in the XynD model. These results offer insight into the highly conserved nature of the catalytic residues, whilst the variation of other residues involved in binding is rather variable and may potentially play a role in enzyme specificity. Trp-219 is found exclusively in Gal*f*ase clusters (Fig. S2), whilst the majority of those in the Ara*f*ase clusters contain a Thr-219. This suggests a key role played by the residue in determining substrate specificity.

Analysis of the protein surface, pore size and accessibility to previously known subsites further confirms a suspected galactofuranosidase specificity of XynD. When superimposed against GH43 arabinoxylan arabinofuranohydrolase from BsAXH-m2,3 with bound xylotetraose (3C7G) (Fig. 3A) a loop on XynD appears to obstruct the I and II sites resulting in a far smaller and shallower binding cleft than BsAXH-m2,3 and suggests XynD is not able to bind and process larger arabinoxylan or derived oligosaccharides (Fig. 3B). Further comparison with Arb43A (1GYE) (Fig. 3C) again shows a far less open binding pocket for XynD when superimposed (Fig. 3D).

**Figure 3:**
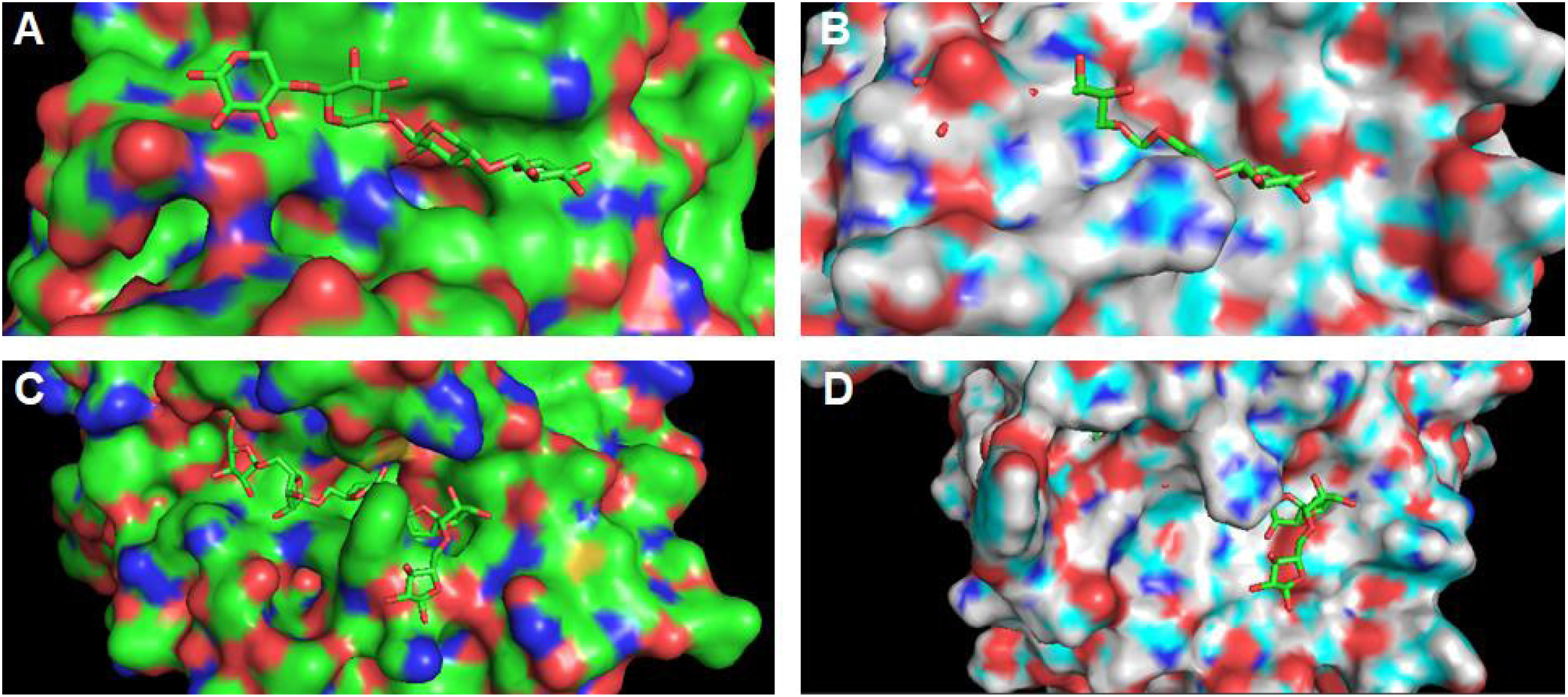
**a** Xylotetraose (X4) binding to *B. subtilis* arabinoxylan arabinofuranohydrolase BsAXH-m2,3 (PDB 3C7G) **b** Superimposition of XynD over 3C7G with X4 ligand **c** Arabinohexaose (A6) binding to *C. japonicus* alpha-L-arabinase Abf43A (PDB 1GYE) **d** Superimposition of XynD over 1GYE with A6 ligand.

### Cloning, expression and purification of XynD

To confirm the activity of XynD experimentally, we heterologous expressed the protein. The predicted signal peptide (residues 1-24) is consistent with transcriptomics data^43^ suggesting that XynD is a secreted protein and that XynD contains four putative N-glycosylation sites. We therefore attempted expression in *Pichia pastoris*, based on a codon optimised cDNA sequence and the pPICZα expression system, but initial trails were unsuccessful. Expression was achieved in *Escherichia coli* Rosetta (DE3) as C-terminally His-tagged XynD without its signal peptide from a pET21a plasmid. The protein was purified via Ni-NTA resin affinity chromatography followed by anion exchange chromatography with a final yield of 7.7 mg l^−1^ culture. Analysis of the purified protein on SDS-PAGE (Fig. S3) indicates that the recombinant enzyme possesses an apparent mass of approximately 35 kDa, which is consistent with theoretical molecular weight of 36.3 kDa.

### Substrate specificity

The activity of XynD was assessed against a panel of ten 4-nitrophenyl-conjugated saccharides. After incubation of 0.5 mg XynD with 1 mM substrate at 25 °C for 2 h, activity was observed only against pNP-β-Gal*f*, while no hydrolysis was detected of pNP-β-Gal*p*, pNP-β-(1,5)-Gal*f*_3_, pNP-α-Ara*f*, pNP-β-Glc, pNP-α-Glc, pNP-β-Xyl, pNP-β-Xyl_2_, pNP-β-GlcNAc, pNP-βlactose. This analysis confirms that XynD is a galactofuranosidase.

Monosaccharide analysis of XynD incubation with galactomannan derived from *A. niger* strain N402, via HPAEC-PAD, detected no release of Gal*f* and thus suggests XynD is not directly involved in degradation of intact galactomannan (Fig. S4). This was confirmed via analysis of the presence of the Gal*f* epitope in galactomannan via the Platelia Aspergillus galactomannan ELISA assay, the abundance of Gal*f* was identical before and after treatment of galactomannan with XynD.

To further test the substrate specificity of XynD, A variety of potential substrates were tested. No XynD activity was detected against the β-(1,5)-linked oligosaccharides 4MU-Gal*f*_3_, 4MU-Gal*f*_4_, 4MU-Gal*f*_5_, 4MU-Gal*f*_6_, a gift of the Oka group^44^. Incubation of XynD with Octyl β-D-Galactofuranosyl-(1,6)-β-D-galactofuranosyl-(1,5)-β-D-galactofuranoside and Octyl β-D-Galactofuranosyl-(1,5)-β-D-galactofuranosyl-(1,6)-β-D-galactofuranoside, a gift from the Lowary group^3^ revealed no observable activity (by MALDI-ToF MS and TLC analysis) against these trisaccharides (Fig. S5). Together these results indicate that XynD is not active on β-1,5- or β-1,6-linked Gal*f* residues.

N-linked glycans on *A. niger* proteins have been described to contain terminal α-(1,2)-linked Gal*f* moities^45^ as well as β-linked Gal*f residues*. To assess activity of XynD on such oligosaccharides, N-linked glycans (Fig. S6) were isolated from Transglucosidase L “Amano” (Amano) as described previously^45^ and purified, and incubated with XynD. Analysis by MALDI-TOF MS (Fig. S7) revealed no observable XynD activity against the glycans. However, as the glycans consisted of a range of structures with mass differences corresponding to one pyranose residue, assessement of activity was not straight forward. The glycan pool was therefore co-digested with jack bean α-mannosidase which cleaves terminal α-(1,2), α-(1,3) and α-(1,6)-mannoses, thus simplifying the glycan structures. A main product structure with a mass of 1095 was detected, corresponding with the mass expected for the sodiated retainment of Gal*f-*α-(1,2)-Man-α-(1,3)-Man-α-(1,6)-Man-β-(1,4)-GlcNAc-β-(1,4)-GlcNAc, thus confirming that glycan structures terminated with Gal*f* (or potentially other non-mannose residues) were obtained. However, no reaction products were observed after incubation of this substrate with XynD, confirming XynD is not active on these Gal*f* N-glycans.

We conclude that the natural substrate of XynD may either be a disaccharide or consist of a glycan with a different linkage type from β-(1,5), β-(1,6) or α-(1,2), for example such as β-(1,2)-linked Gal*f* as found in glycoinositolphospholipids.

### Biochemical characterization

Using Gal*f*-pNP as a model substrate, we assessed the optimum reaction conditions and kinetic parameters for XynD. The enzyme exhibited its highest activity at pH 4-5, as assessed after incubation of purified XynD at 25 °C for 1 h in a range of citrate-phosphate buffers at pH 3-9 (Fig. 4A). At pH >5 the enzyme showed substantial reduction in activity. This is similar to a previously described *A. niger* β-D-galactofuranosidase that demonstrated optimal activity at pH 3-4^32^. XynD temperature optimum was 25 °C (Fig. 4B) with substantial decrease in activity observed after incubation at temperatures greater than 30 °C. The temperature stability of XynD decreased when incubated above 30 °C for 1 h (Fig. S8).

**Figure 4:**
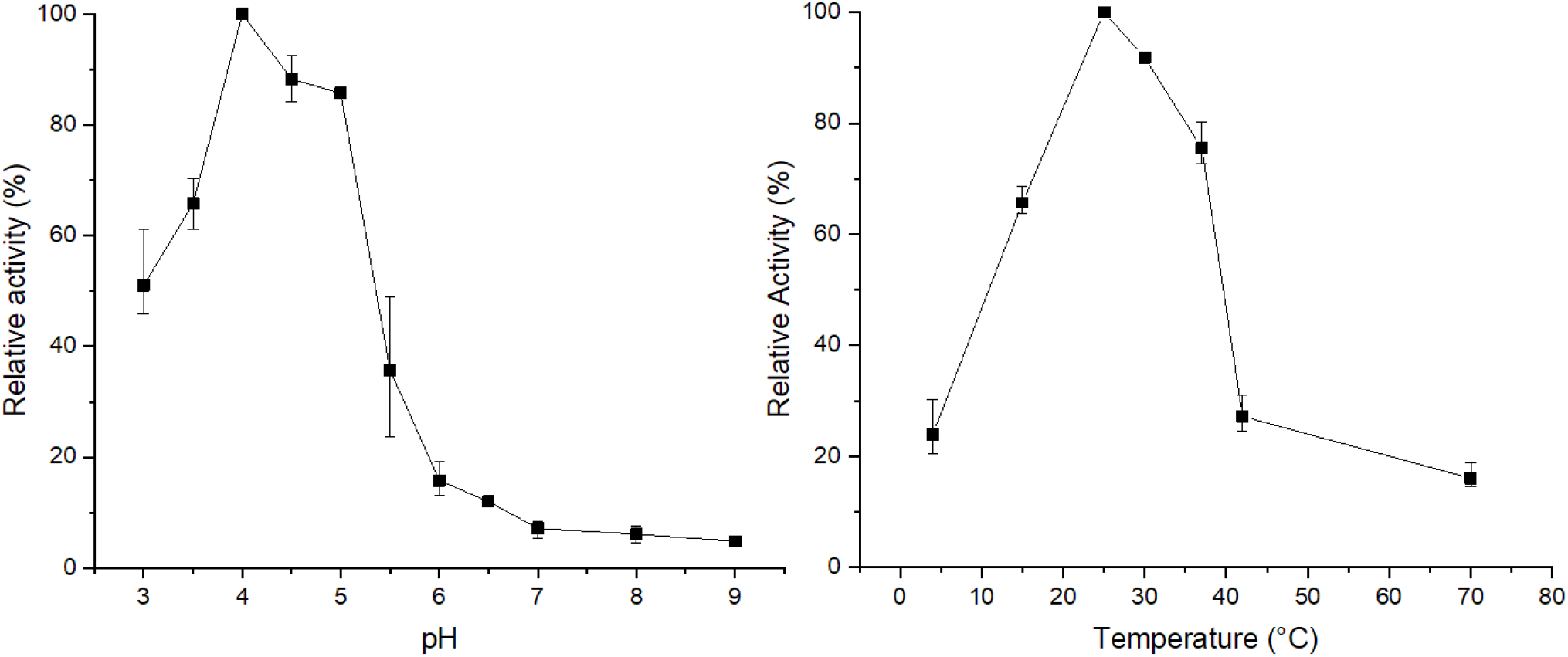
Optimisation of reaction conditions for pNP-Gal*f* hydrolysis by XynD **a** Relative activity at pH 3-9 **b** Relative activity at tempera-tures 4-70 *°C*

The specific activity of XynD against *p*NP-β-Gal*f* at 25 °C, pH 5 was 0.23 μmol min^−1^ mg^−1^. The activity appears to follow Mi-chaelis-Menten kinetics, displaying a K_m_ of 17.9 ± 1.9 mM, V_max_ of 70.6 ± 5.3 μM min^−1^ and K_cat_ of 0.14 s^−1^.

## Discussion

Gal*f* containing glycans such as galactomannan or O-glycans are important as drug targets and immunogenic motifs in the production of pharmaceuticals due to the absence of the Gal*f* monosaccharide in mammalian tissues^23^. There has therefore been a longstanding interest in elucidation of the enzymatic mechanism contributing to the turnover of such glycans^1^.

Galactomannan degrading activity of an unidentified enzyme produced by *A. niger* has been previously reported^32^. Subsequent investigation to identify this source of activity has revealed a variety of Gal*f* active enzymes within the *Aspergillus* genus. *A. niger* AbfA and AbfB both have dual activity against pNP-Gal*f* and pNP-Ara*f* yet do not degrade galactomannan^33^. More recently, *A. nidulans* exo-acting Gal-*f*ases GfgA and GfgB have been identified, and have been shown to hydrolyze both β-(1,5) and β-(1,6) linkages, whilst *A. fumigatus* Afu2g14520 demonstrates dual activity against pNP-Gal*f* and pNP-Ara*f,* with higher activity against the latter^34^. Here we demonstrated that XynD encodes an galactofuranosidase active against pNP-Gal*f.* The enzyme is not responsible for galactomannan degradation and lacks activity against β-(1,5), β-(1,6)- and α-(1,2) linked Gal*f* substrates. Instead its activity suggests glycoinositolphospholipids may be the natural substrate.

XynD has previously been annotated as a putative xylosidase in the literature and our identification of its galactofuranosidase activity highlights how detailed phylogenetic analysis combined with biochemical characterization is critical to understand the roles of glycoside hydrolases. Our phylogenetic analysis of the subfamily, followed by mapping of biochemically characterized members, demonstrated how GH43-34 is grouped into multiple clusters containing characterized members with arabinofuranosidase activity. Assignment of Cluster I members as arabinofuranosidases was further supported by association of a subset of members from this cluster with CBM42, which are commonly associated with GH54 arabinofuranosidases and bind arabinofuranose present in arabinoxylan^46^. Only cluster II contained bacterial members with galactofuranosidase activity and our structural modelling, substrate docking and biochemical characterization confirmed our hypothesis that as fungal member of this cluster, XynD, had galactofuranose activity.

Previously studies of GH family 43 reveal only the catalytic residues (Asp, Glu, and its p*K*_a_-modulating Asp) are well conserved^42^, and our molecular docking studies confirm the binding pose of galactofuranose in XynD is stabilized to these from local Asp, Glu, Trp and Arg residues. In XynD, many potential subsites are completely occluded by surrounding loops between separate beta-strands and thus explains the lack of catalysis demonstrated against large oligo- and polysaccharides. The variety of specificities observed in the GH43 family stem from the variable orientation of which the substrate can sit within the deep pocket of the active site, thus resulting in a diverse and complex family^47,48^.

Our modelling of the XynD structure combined with sequence alignments highlighted how an amino acid in the active site of the GH43-34, is consistently different between the arabinofuranosidases and galactofuranosidases and could affect the substrate specificity in the GH43-34 subfamily. This W219, whose equivalent is a T in arabinofuranosidases, is positioned to affect sugar binding in the subsite and we hypothesised that mutation of this site may switch enzyme activity from a galactofuranosidase to and arabino-furanosidase. Attempts to a form a W219T mutant of XynD were unsuccessfull due to a lack of protein production from a mutant plasmid.

XynD was the first fungal member of GH family 43-34 to be recombinantly expressed and biochemically characterized. We showed XynD demonstrates hydrolytic activity against pNP-Gal*f.* Interestingly, despite the substantial structural similarities between Gal*f* and Ara*f,* XynD does not show activity against pNP-Ara*f* and is thus thereby differs from other characterized *A. niger* galactofuranosidases that exhibit dual activity^33^. An unidentified *A. niger* enzyme with Gal*f*ase activity has been studied^32^. The enzyme is substantially larger than XynD at ~80 kDa (after N-glycan removal by EndoH treatment), although this difference could be accounted for by the presence of O-linked glycans. The unidentifed Gal*f*ase exhibits a lower a K_m_ (4 mM) in comparison to that of XynD (17.9 mM), and therefore likely is a different enzyme from XynD.

XynD displays an optimum pH between pH 4 and pH 5, and concurs with previously described recombinant galactofuranosidases with varying optimum pHs of 3-7^28,30,34,49,50^. Optimal temperature of characterized galactofuranosidases vary, for example, the Galf-ase derived from Streptomyces sp. JHA26 lost its activity when incubated at a temperature over 45 °C whilst Streptomyces sp. JHA19 galactofuranosidases had an optimal temperature at 60 °C^50,51^. Since XynD was expressed in *E. coli* no post translational modifications, such as glycosylation, will have occurred. Therefore, lack of post translational modifications should be considered when evaluating parameters such as activity, pH optimum and temperature optimum.

To gain more insight in the biological role of XynD, we assessed the conditions under which *xynD* expression and XynD secretion has been reported. Gene co-expressing networks, based on 155 transcriptomics experiments^52^ and available via FungiDB, highlighted expression of *xynD* is positively correlated with 110 genes, and gene ontology (GO) enrichment analysis using REViGO^53^ (Fig. S9) revealed that these are enriched for transmembrane transporter activity (GO:0022857), oxidoreductase activity (GO:0016491), hydrolase activity, acting on glycosyl bonds (GO:0016798) and hydrolase activity with activity on carbon-nitrogen (but not peptide) bonds (GO:0016810). This positive correlation suggest *xynD* is expressed during carbohydrate degradation. In *A. niger*, *xynD* expression has previously been reported to be induced in response to plant-derived polysaccharides and lignocellulose, despite lack of putative binding sites in its promoter for the master regulator XlnR. Low levels of expression *xynD* are observed on a variety of easily metabolised carbon sources (glucose, sucrose, fructose and sugar beet pulp) and expression is suspected to be subject to carbon catabolite repression, as supported by 11 binding sites for carbon catabolite repressor CreA in the promotor region of *xynD*^54^ and other genes undergoing such repression^55^. Whilst *xynD* is not expressed during growth on maltose in the fungal wild-type^55^, mutants with an inactivated *amyR* (encoding a regulator of starch degradation induced by maltose^56^) display upregulation of *xynD* transcription^55^. Similarly, *xynD* expression on arabinose is low in the wild-type, but an *araR* disruption mutant showed upregulation of *xynD* transcription^55^. In the *amyR*^57^ and *araR*^58^ mutants the expression of sugar transporters and metabolic enzymes that enable growth on maltose and arabinose, respectively, will be negatively affected and thus the observed expression pattern is consistent with a response to carbon limitation and concurs with the expression response to starvation in other *A. niger* galactofuranosidases^32^.

In conclusion, this study revealed further insight into the degradation of galactofuranose containing biomolecules with the identification of a GH43-34 *A. niger* galactofuranosidase. Understanding this vast but largely undescribed subfamily could play a key role in developing knowledge of glycosylation in a variety of fungal products, such as pharmaceuticals.

## Experimental procedures

### Phylogenetic analysis of XynD

GH 43-34 domains were identified using the Pfam data base^59^ then subsequently aligned and trimmed to just the catalytic domain using BioEdit^60^. Evolutionary analyses were conducted in MEGA X^61^. The evolutionary history was inferred by using the Maximum Likelihood method and JTT matrix-based model^62^. Initial tree(s) for the heuristic search were obtained automatically by applying Neighbour-Join and BioNJ algorithms to a matrix of pairwise distances estimated using the JTT model, and then selecting the topology with superior log likelihood value. The tree was drawn to scale, with branch lengths measured in the number of substitutions per site. This analysis involved 963 amino acid sequences. There were a total of 494 positions in the final dataset.

### Computational docking studies

Enzyme modelling and molecular docking simulation was performed using the YASARA software from YASARA Biosciences GmbH. As there is no PDB structure of XynD a model was produced using *Bacteroides thetaiotaomicron* endo-1,4-beta-xylanase D (PDB 3QZ4) as a template and was subsequently used to generate the receptor for simulations. In all cases, the raw crystal structures were first prepared for simulation using the YASARA “Clean” script, which detects and amends crystallographic artefacts and assigns protonation states. Before docking, the energy minimization script of the YASARA software package was applied to the protein in order to obtain the most likely starting structure. For energy minimization, a simulation cell was defined around the whole protein and the cell was filled with water molecules. For docking, the water was removed afterwards with a variety of arabinose and galactofuranose based docking ligands. Ligands were minimised in vacuo using the energy minimization script of the YASARA software package. The docking simulation itself was performed using the dock_run.mcr macro using VINA as the docking method and the AMBER03 force field. Appropriate simulation cells were defined for the respective docking simulations. The created receptor-ligand complex structures were further processed using the PyMOL software from Schrodinger LLC involving the identification of polar contacts between the ligand and the receptor, as well as determination of bond distances.

### Construction of expression vectors

The gene *xynD* (An11g03120, GenBank: CAK40644.1), with an 86 bp intron removed, was synthesised by GeneArt gene synthesis (Thermo Fisher scientific) with codon optimisation for expression in *Pichia pastoris*. Construct included 5’ and 3’ complementary overhangs for ligation into pPICZαA using NEBuilder HiFi DNA Assembly (New England Biolabs) to form pPICZαA_GH43_34. The coding sequence of *xynD* gene was amplified from pPICZαA_GH43_34 using primers 5’ CGC<u>CATATG</u>CACCCACAACAGAACTTG-3’ (forward) and 5’-CGC<u>CTCGAG</u>AGACAAAGTTCTACCCTCGACAC-3’ (reverse). The primers contained restrictions sites *Nde*I and *Xho*I (underlined) for forward and reverse primers, respectively. The PCR conditions were as follows: initial denaturation 95 °C (5 min), [95 °C (30 s) denaturation, 58 °C (30 s) annealing and 72 °C (30 s) extension] for 20 cycles, final extension 5 min. PCR amplifications were conducted using Phusion DNA polymerase. The resulting products were run on 1 % agarose gel then extracted using QIAquick Gel Extraction Kit (Qiagen). Restriction enzymes and Phusion DNA polymerase were purchased from New England BioLabs Inc and used according to the manufacturer’s instructions. The amplified fragment was cloned into linearized (NdeI and XhoI) pET21a to form pET21a_43_34. Correct insertion of the gene in the expression vector was confirmed by sequencing. The amino acid sequence of the recombinant XynD is as follows: MH-PQQNLLATTTSNTKAGNPVFPGWYADPEARLFNAQYWIYPTYSA-DYSEQTFFDAFSSPDLLTWTKHPTILNITNIPWSTNRAAWAPS-VGRKLRSSANAEEEYDYFMYFSVGDGTGIGVAKSTTGKPEGPY-EDVLGEPLVNGTVYGAEAIDAQIFQDDDGRNWLYFGGWSHAV-VVELGEDMISLKGDYLEITPEGYVEGPWMLKRNGIYYYMFSVG-GWGDNSYGVSYVTADSPTGPFSSTPKKILQGNDAVGTSTGHNS-VFTPDGQDYYIVYHRRYVNDTARDHRVTCIDRMYFNEAGEILPVNITLEGVEGRTLS(LEHHHHHH-)

### Protein expression and purification

*E. coli* strain Rosetta™(DE3) carrying pET21a_43_34 was in-oculated into LB media supplemented with 100 ug ml^−1^ ampicillin and grown at 37 °C, 250 rpm until OD_600_ ~0.25. Cells were then induced with 1mM IPTG final concentration and grown at 22 °C, 250 rpm for 16 h. Cells were harvested by centrifugation (4000 rpm, 15 min, 4 °C), suspended in His-buffer (Tris 50 mM, NaCl 250 mM, imidazole 20 mM, pH 8) and lysed by sonication (20s on, 20s off, 5 min). Cell debris was removed by centrifugation (4000 rpm, 15 min, 4 °C). Protein was purified by metal affinity chromatography on a Ni^2+^-NTA column (Qiagen). Cell free extract was loaded onto column followed by two wash steps with wash buffer (Tris 50 mM, NaCl 250 mM, imidazole 20 mM, pH 8) followed by application of elution buffer (Tris 50 mM, NaCl 250 mM, imidazole 200 mM, pH 8), all fractions were retained for SDS-PAGE analysis. The his-tagged purified protein was then further purified using anion exchange chromatography employing a HITrapQ HP column 1 mL, whereby the protein was eluted with a gradient of 20 mM Tris-HCl buffer, 0.5M NaCl, pH 7.2. Eluted proteins were analysed by 4—15% sodium dodecyl sulfate polyacrylamide (SDS-PAGE) gel electrophoresis (Mini-PROTEAN® TGX™ Precast Gels by Bio-rad) using broad range molecular weight markers (PageRuler™ Plus Prestained Protein Ladder) as standard, and the protein bands were visualised by staining with Coomassie Brilliant Blue G250. The protein concentration was determined by Bradford method using bovine serum albumin (BSA) as a standard based on the protocol provided by Sigma-Aldrich (96 Well Plate Assay Protocol). All experiments presented in this manuscript were performed with XynD purified by anion exchange chromatography unless otherwise stated.

### Mass spectrometry of carbohydrates

Samples were prepared for analysis by MALDI-TOF via crystallisation with super DHB or THAP matrices. Super DHB (a 9:1 (w/w) mixture of 2,5-Dihydroxybenzoic acid and 2-hydroxy-5-methoxybenzoic acid) was prepared at 15 mg ml^−1^ in a mixture of 50 % (v/v) acetonitrile and 50 % (v/v) water containing 0.1 % trifluoroacetic acid (TFA). 2′,4′,6′-Tri-hydroxyacetophenone monohydrate (THAP) was prepared at 10 mg ml^−1^ in acetone. MALDI-TOF mass spectrometry was performed using the Bruker Ultraflex 3 in positive mode. MALDI-TOF was calibrated using peptide Calibration Standard II (Bruker) containing a range of peptides 757-3149 Da in size.

### Substrate specificity

XynD (0.5 mg ml^−1^) was incubated overnight at 25 °C in 50 mM citric acid-phosphate pH 5 with pNP-labelled substrates at 1 mM. All experiments were performed in triplicate. Reactions were halted via a 1:1 volume addition of Na_2_CO_3_ 1M with pNP release measured at OD_405_. The following substrates were used: pNP-β-D-galactofuranoside (pNP-β-D-Gal*f*; Carbosynth), pNP-α-ʟ-arabinofuranoside (pNP-α-ʟ-Ara*f*; Megazyme), pNP-β-galactopyranoside (pNP-β-Gal*p*; Sigma-Aldrich), pNP-β-glucopyranoside (pNP-β-Glc*p*; Sigma-Aldrich), pNP-α-glucopyranoside (pNP-α-Glc*p*; Sigma-Aldrich), pNP-N-acetyl-β-D-glucosaminide (pNP-β-D-GlcNAc; Sigma-Aldrich), pNP-lactose (pNP-Lac; Sigma-Aldrich), pNP-xylopyranoside (pNP-Xyl*p*; Sigma-Aldrich), pNP-xylobiose (pNP-β-Xyl_2_, Carbosynth), pNP-β-(1,5)-Gal*f-*β-(1,5)-Gal*f*-β-(1,5)-Gal*f* (pNP-β-(1,5)-Gal*f*_3_); gift of Dr. T. Oka.

XynD (0.5 mg ml^−1^) was incubated overnight at 25 °C in 50 mM citric acid-phosphate pH 5 with the following 4MU-labelled substrates at 1 mM: 4MU-Gal*f*_3_, 4MU-Gal*f*_4_, 4MU-Gal*f*_5_, 4MU-Gal*f*_6_, (gifts of Dr. T. Oka, Sojo University, Ja-pan). Reactions were halted by the addition of stop solution (1M sodium hydroxide, 1M glycine, pH 10) with released 4-MU detected after excitation at 365 nm and emission at 440 nm.

XynD (0.5 mg ml^−1^) was incubated overnight with 1 mM final concentration Octyl β-D-Galactofuranosyl-(1,6)-β-D-galactofuranosyl-(1,5)-β-D-galactofuranoside and Octyl β-D-Galactofuranosyl-(1,5)-β-D-galactofuranosyl-(1,6)-β-D-galactofuranoside (gifts of Dr. T.L. Lowary, The Institute of Biological Chemistry, Academia Sinica, Taiwan) in 50 mM citric acid-phosphate buffer. Reaction were then monitored by MALDI-TOF MS.

The α-(1,2)-Gal*f*-containing N-linked glycans were released from 1 mg Transglucosidase L “Amano” (Amano) by using PNGase F (New England Biolabs) under denaturing conditions. Reactions were then filtered via Vivaspin 500 (Sigma-Aldrich) to obtain the glycan products. Samples were then purified using a 1 ml Supelclean™ ENVI-Carb™ SPE Tube. The column was prepared with 1 ml of each solution in a sequential order: 75% acetonitrile + 0.1% trifluoroacetic, 1 M NaOH, water, 30% acetic acid, water, 75% acetonitrile + 0.1% trifluoroacetic. The sample was then applied and washed with 3 column volumes of water followed by subsequent elution with 3 mL 25% acetonitrile + 0.1 % trifluoroacetic acid. The N-linked glycans were then dried and resuspended in water. Glycans were incubated with final concentation 1 mg ml^−1^ XynD in 50 mM citric acid-phosphate buffer, pH 5 overnight, then analysed by MALDI-TOF MS. Additionally glycans were co-incubated with 1 mg ml^−1^ XynD and 1 mg ml^−1^ α-Mannosidase from *Canavalia ensiformis* (Sigma-Aldrich).

Activity of XynD on *A. niger* galactomannan, a kind gift from Dr A. Ram and J. Park (The University of Leiden, the Netherlands), was assessed via overnight incubations containing final concentration 0.5 mg ml^−1^ XynD and 1 mg ml^−1^ *A. niger* N402 galactomannan in 50 mM citric acid-phosphate buffer at 25 °C. Reactions were then filtered via Vivaspin 500 (Sigma-Aldrich) to remove protein from the reactions. Galactomannan, hydrolysed via incubation in trifluoroacetic acid at 100 °C for 5 h acted as a positive control for detection of constituant monosaccharides XynD activity was assessed via detection of released galactose by HPAEC-PAD as described in detail below, and via Platelia™ Aspergillus Ag (Bio-Rad), as detailed by the manufacturer.

### HPAEC-PAD analysis

High performance anion exchange chromatography (HPAEC) was performed to analyze monosaccharide release by XynD, based on a published method^63^. A Thermo Scientific Dionex ICS-6000 HPAEC system with pulse amperometric detection (PAD) controlled by Chromeleon software version 7.1 was used. A CarboPac PA1 guard (2 mm x 50 mm, Thermo Fisher Scientific) and a CarboPac PA1 analytical column (250 mm × 2 mm, Thermo Fisher Scientific) for monosaccharide analysis were employed. Monosaccharides were eluted isocratically using H_2_O and 1M NaOH in a ratio of 80:20 for 30 min with a flow rate of 0.25 ml min^−1^.

### Identification of pH and temperature optimum and stability

To identify the optimal pH for activity, 1 mM pNP-β-Galf was incubated with XynD (0.5 mg ml-1) for 1 h at 25 °C in 50 mM citric acid-phosphate and Tris-HCl buffers at a range of pH 4-9. Reactions were halted via a 1:1 volume addition of 1M NaCO3, after which pNP release as measured at OD_405_. To identify the optimal reaction temperature, reactions were incubated in 50 mM pH 5 citric acid-phosphate buffer at a range of temperatures from 4-90 °C. All experiments performed in triplicate and enzyme activity was linear over time under the conditions described. pH stability was determined via pre-incubation of the enzyme (0.3 mg ml^−1^) for 1 hat pH 3-9 before assaying residual activity on pNP-Galf under standard condition reactions at pH 5.

### Co-expression analysis of XynD

*A. niger* microarray data from FungiDB was used to elucidate co-expression networks. Spearman scores of 0.5 of greater are considered significantly co-expressed. 1241 / 110 genes showed a positive correlation with the *xynD* gene (Spearman coefficient ≥ 0.5 / ≥ 0.75) and 1476 / 70 genes a negative correlation (Spearman coefficient ≥−0.5 / ≥−0.75).

## Acknowledgements

We thank Dr A. Ram and J. Park of The University of Leiden, the Netherlands, for providing us with *A. niger* galactomannan, Dr T.L Lowary of The Institute of Biological Chemistry, Academia Sinica, Taiwan for providing us with the (1,5),(1,6) and (1,6),(1,5) galactofuranoside trisaccharides and Professor T. Oka of Sojo University, Japan, for the pNP and 4-MU based galactofuranosides. We are grateful to Dr W. Finnigan and R. Sung of The University of Manchester, UK for their assistance with bioinformatics and HPAEC-PAD analysis, respectively.

## Funding and additional information

This study was funded by the BBSRC via BB/P011462/1 (JvM), the EPSRC via DTP EP/N509565/1 supporting GB, an undergraduate bursary from the British Mycological Society supporting FWY and a Learning Through Research Internship from the University of Manchester supporting NB.. BSJ and SLF acknowledge funding by the European Research Council under grant agreement no. 788231-ProgrES-ERC-2017-ADG.

