## supplementary data for "Biochemical characterization of a glycoside hydrolase family 43 β-D-galactofuranosidase from the fungus *Aspergillus niger*"

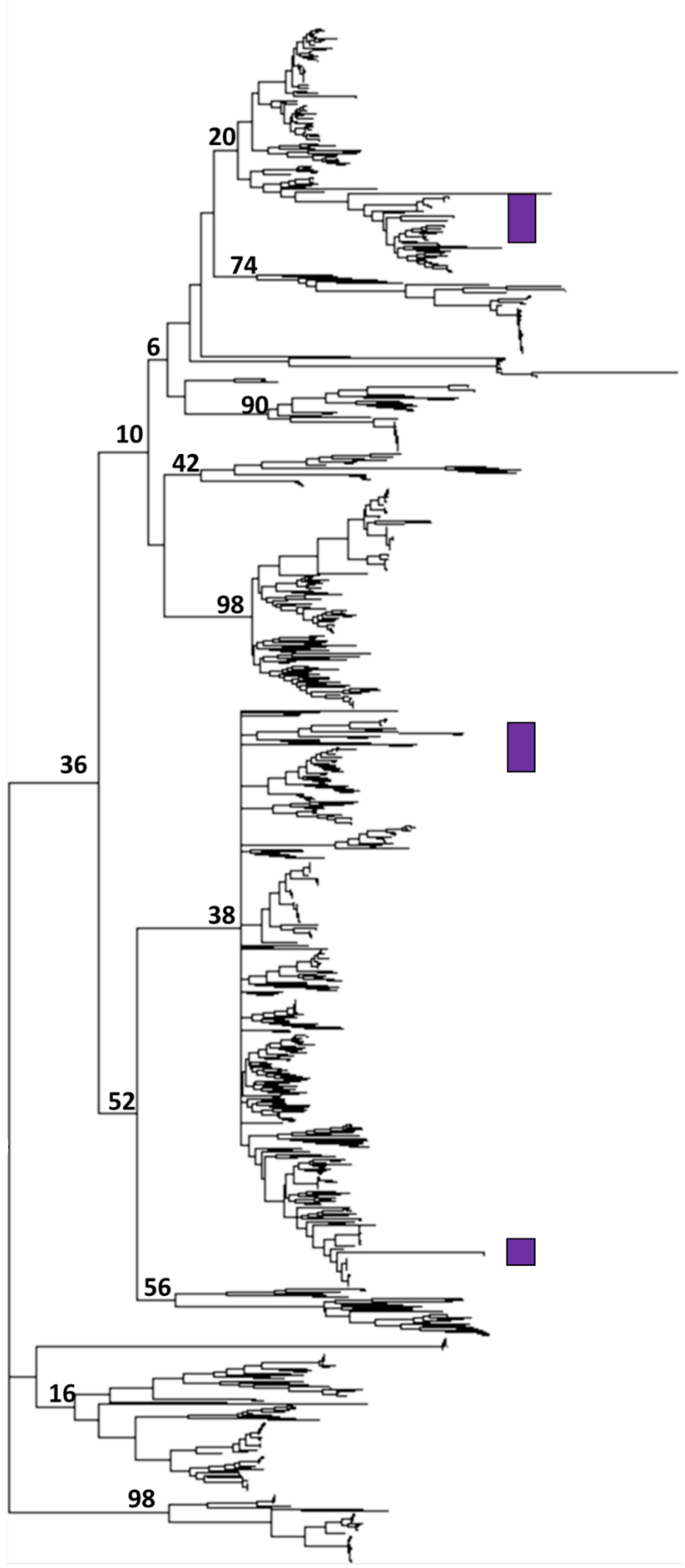


**Supplementary figure 1**: Phylogenetic tree of GH43 subfamily 34, annotated with boostrap values and locations of fungal sequences highlighted (purple).

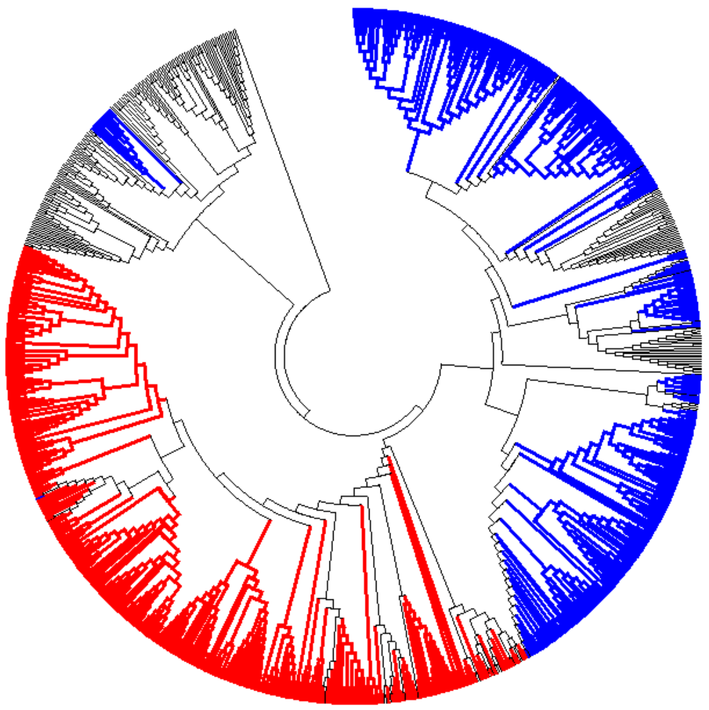

**Supplementary figure 2**: Phylogenetic tree of GH43 subfamily 34 with amino acid identity at position 219 highlighted: W (red), T (blue) or other (grey)

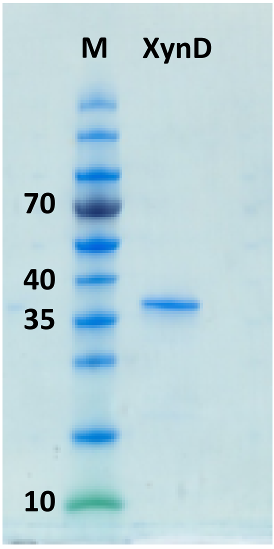


**Supplementary figure 3**: SDS-PAGE analysis of heterologously expressed XynD after His-tag purification. M: Molecular weight Marker.

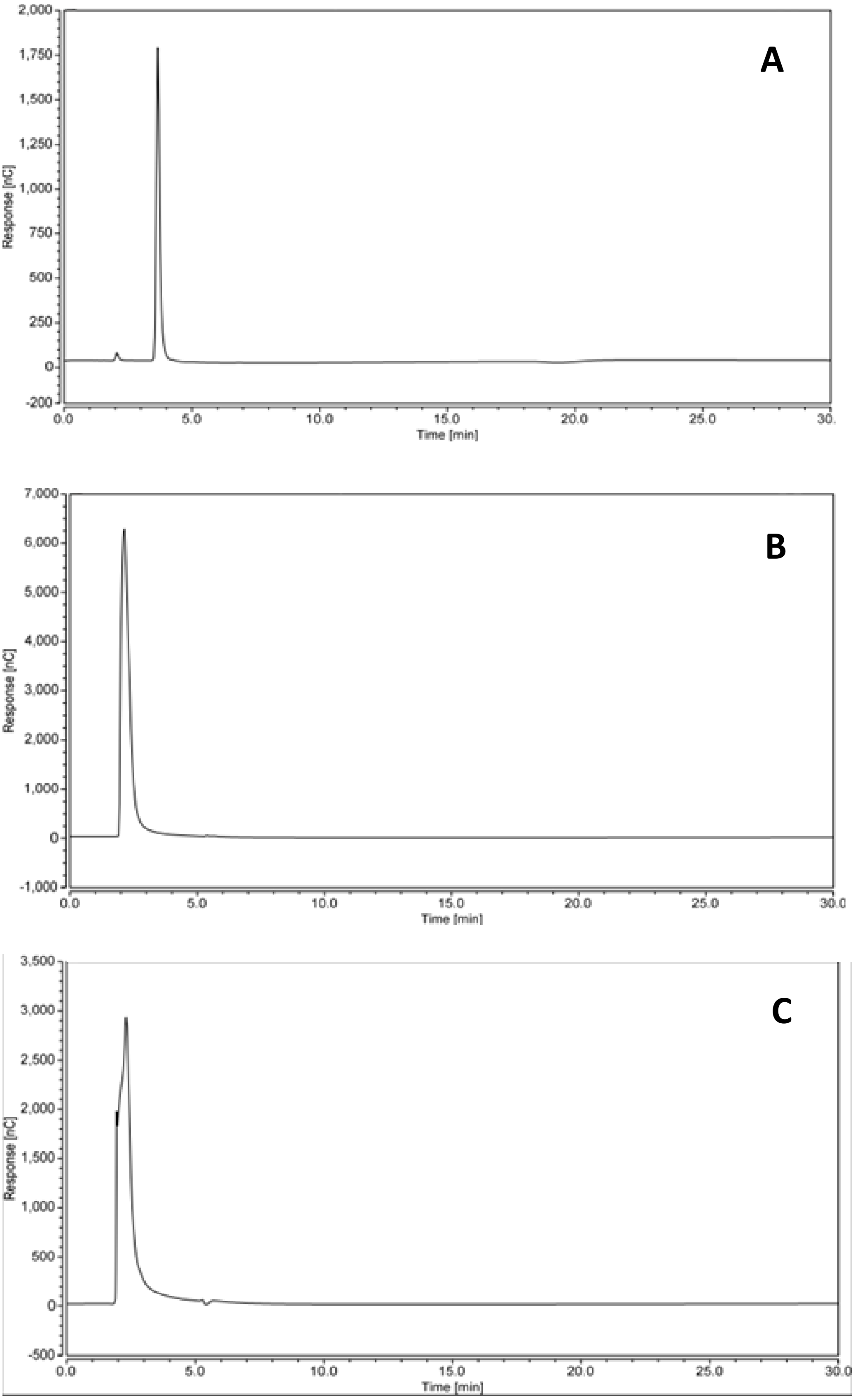

A
Response [nC]
Time [min]
2,000
1,750
1,500
1,250
1,000
750
500
250
0
-200
0.0
5.0
10.0
15.0
20.0
25.0
30.
B
Response [nC]
Time [min]
7,000
6,000
5,000
4,000
3,000
2,000
1,000
0
-1,000
0.0
5.0
10.0
15.0
20.0
25.0
30.0
C
Response [nC]
Time [min]
3,500
3,000
2,500
2,000
1,500
1,000
500
0
-500
0.0
5.0
10.0
15.0
20.0
25.0
30.0

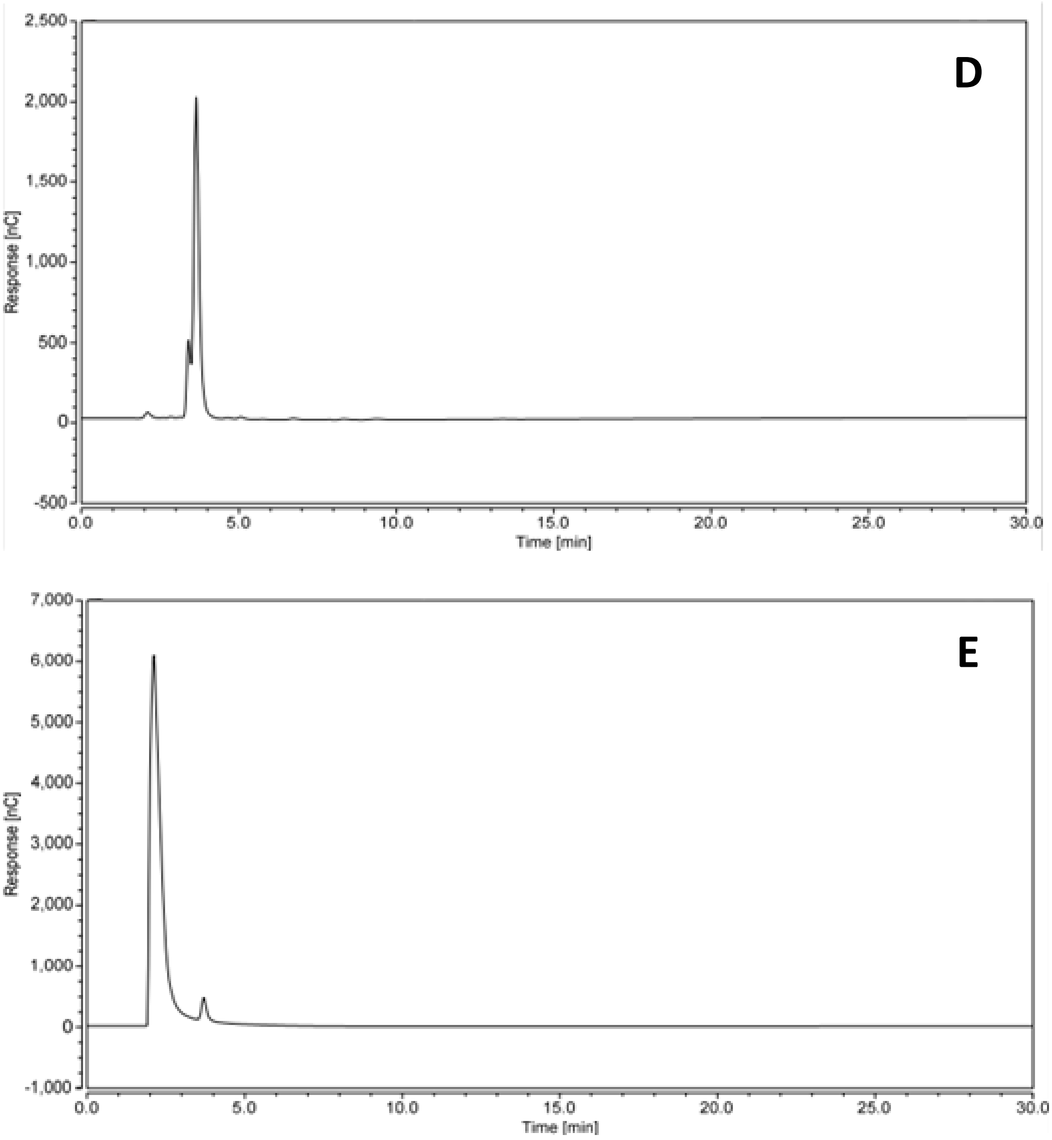


**Supplementary figure 4**: High-Performance Anion Exchange Chromatography (HPAEC) analysis of XynD activity on *A. niger* N402 galactomannan. **A** 1 mM Galactose **B** *A. niger* N402 galactomannan after incubation with XynD **C** *A. niger* N402 galactomannan incubation negative control (incubation with protein fraction as obtained after purification of an *E. coli* culture expressing an empty pET 21a vector under conditions identical to those used to obtain XynD) **D** Acid-hydrolysed *A. niger* N402 galactomannan **E** *A. niger* N402 galactomannan empty vector incubation spiked with galactose

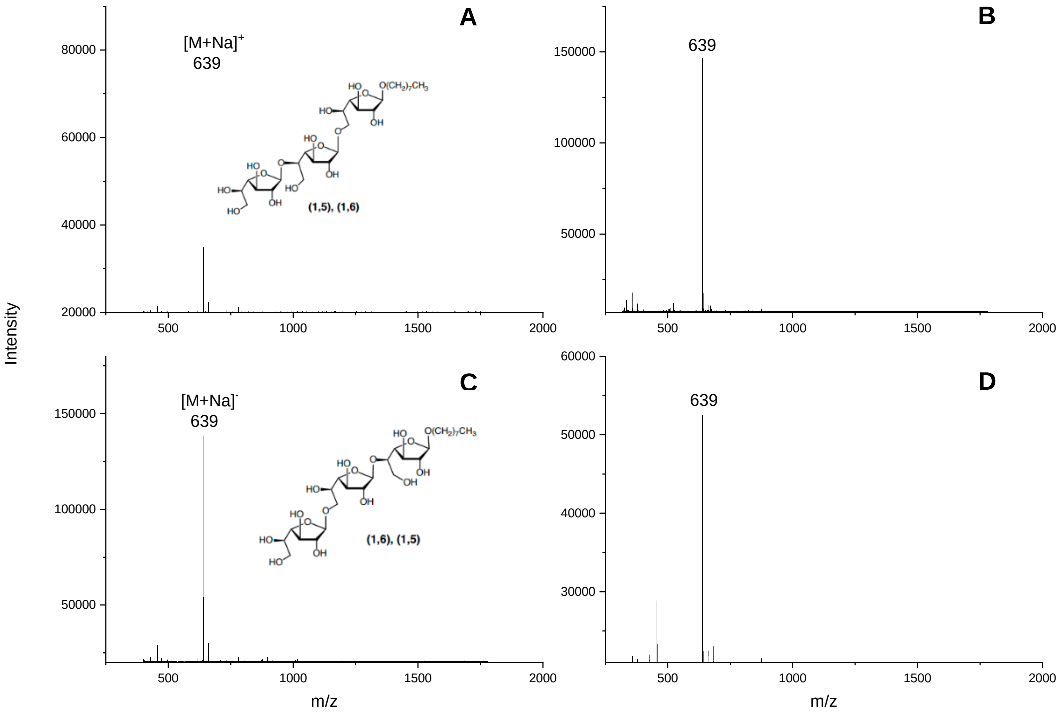


**Supplementary figure 5**: Assessment of XynD activity on β-1,5/β-1,6 Gal*f* trisaccharides via MALDI-TOF MS. **A** Octyl β-D-Galactofuranosyl-(1,5)-β-D-galactofuranosyl-(1,6)-β-D-galactofuranoside (M = 616) **B** Octyl β-D-Galactofuranosyl-(1,5)-β-D-galactofuranosyl-(1,6)-β-D-galactofuranoside after incubation with XynD **C** Octyl β-D-Galactofuranosyl-(1,6)-β-D-galactofuranosyl-(1,5)-β-D-galactofuranoside (M = 616) **D** Octyl β-D-Galactofuranosyl-(1,6)-β-D-galactofuranosyl-(1,5)-β-D-galactofuranoside after incubation with XynD. Expected mass of 477 if Gal*f* residue removal (not observed).

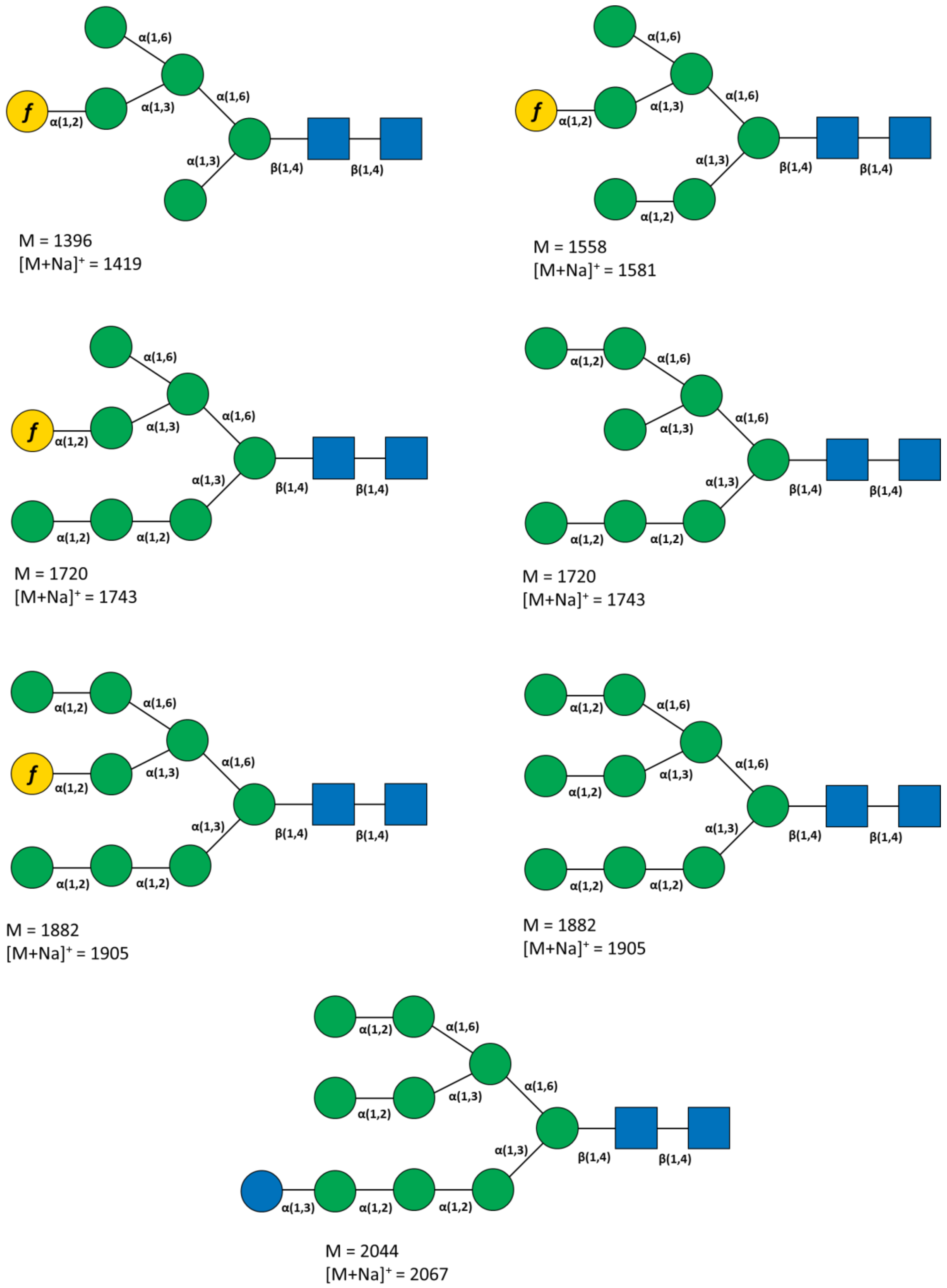


**Supplementary figure 6:** N-linked glycans produced by incubation of Transglucosidase L "Amano" (Amano) with PNGase F, as described in [1].

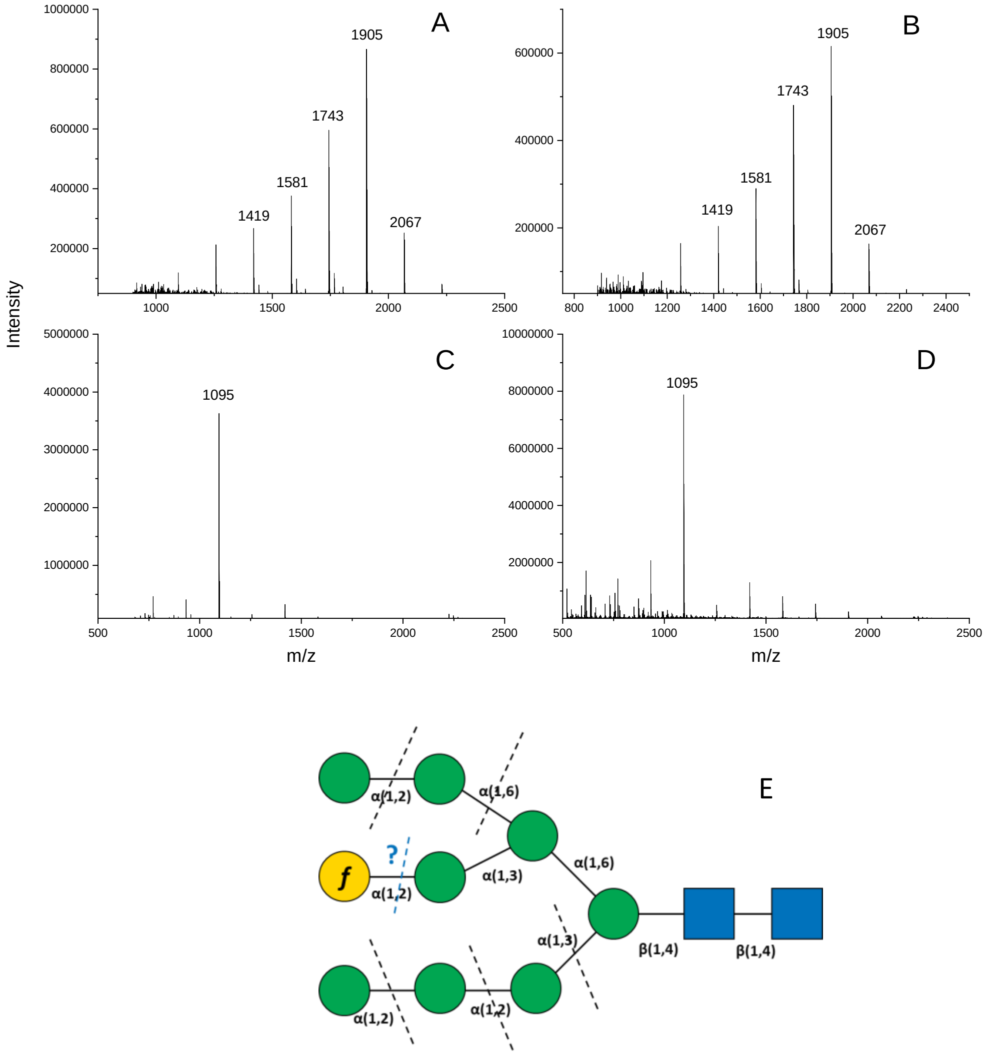


**Supplementary figure 7**: MALDI-TOF MS of α-1,2 containing-N-linked glycans incubated with XynD. **A** Purified Amano Transglucosidase N-linked glycans **B** Purified Amano Transglucosidase N-linked glycans XynD incubation. No shift of -162, consistent with terminal Gal*f* removal was observed. **C** Purified Amano Transglucosidase N-linked glycans co-incubated with mannosidase and XynD **D** Purified Amano Transglucosidase N-linked glycans incubated with mannosidase**. E** Diagram demonstrating enzymatic activity against N-linked glycan, black dashed lines representing exo-mannosidase activity, blue dashed line and question mark representing potential Gal*f*ase activity. Lack of Gal*f*ase activity blocks mannosidase activity against Gal*f*-terminating chain, leaving the hexasaccharide detected in panels C and D. Conversely, Gal*f*ase activity would enable degradation of the glycan to the trisaccharide Man-GlcNAc-GlcNAc core ($[M+Na]^+$= 609).

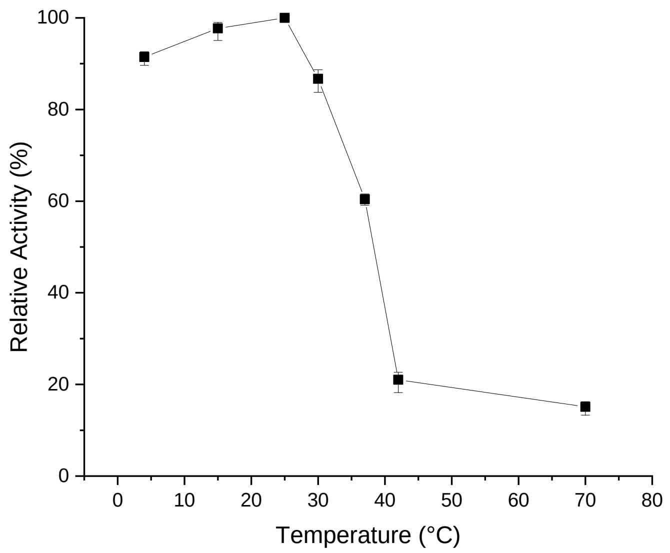


**Supplementary figure 8**: Temperature stability of XynD. After 1 h incubations at indicated temperatures, residual activity of XynD was measured under standard reaction conditions (pH 5, 25 °C).

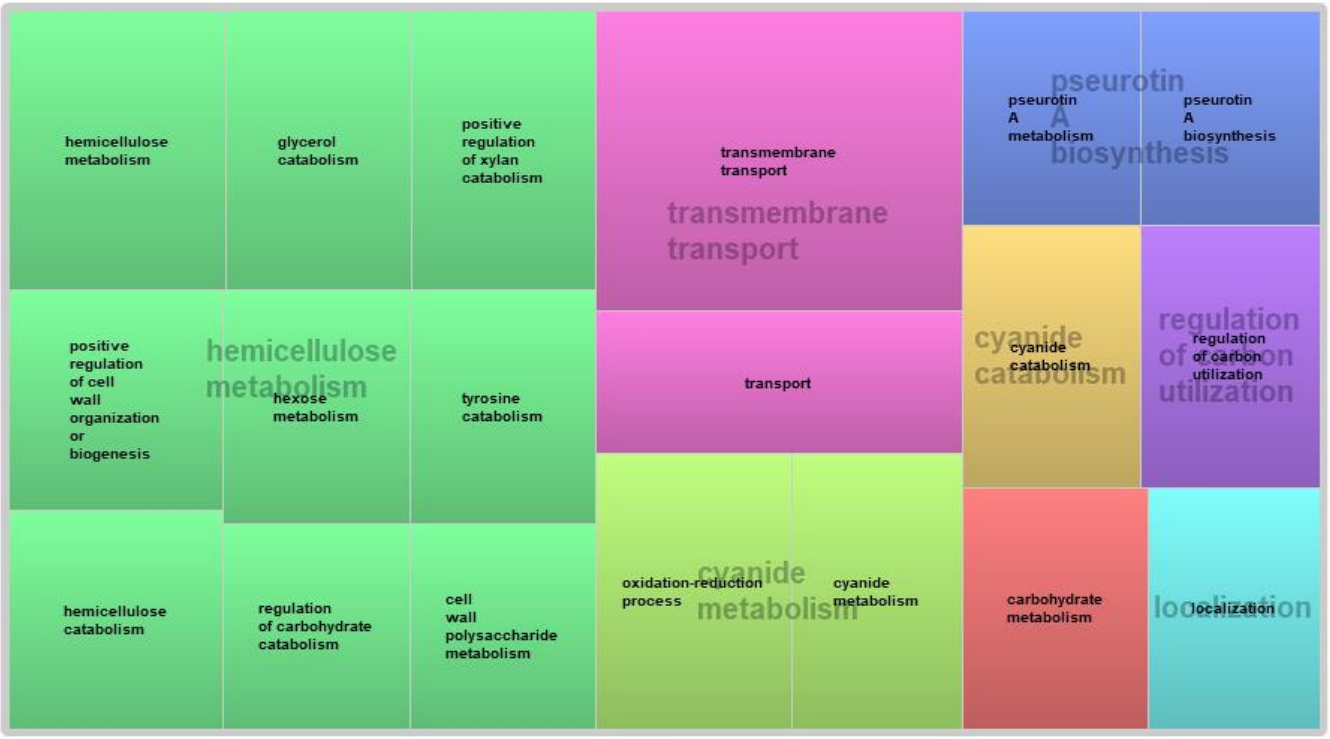


**Supplementary figure 9**: Revigo TreeMap of GO terms (biological processes ontology) positively correlating with *xynD* expression
